# *ggrecipes*: An R Package for Custom Visualizations using *ggplot2*

**DOI:** 10.64898/2025.12.18.694217

**Authors:** Ignophi Hu

## Abstract

We introduce *ggrecipes*, an R package that provides a collection of *ggplot2*-based functions for data exploration and analysis. Each function encapsulates data preprocessing and visualization construction, and returns a *ggplot2* object that can be further customized using standard *ggplot2* syntax. The package includes both general-purpose and domain-specific visualizations. *ggrecipes* is implemented in R under an open-source MIT license, with source code and documentation available at https://github.com/Ignophi/ggrecipes and https://ignophi.github.io/ggrecipes/.

## 1. Introduction

Visual exploration is a central tool for hypothesis generation, model development, and result interpretation. The R computing environment provides a rich ecosystem for custom visualization. Among available tools, ggplot2 [1] has become a widely adopted solution. It implements a layered grammar of graphics [2] that is both flexible and expressive. This approach enables the construction of domain-specific visual representations that arise from specific questions. As similar questions recur, certain visual patterns prove repeatedly useful, becoming part of a field’s analytical vocabulary.

Custom visualizations generally follow two steps. First, data are preprocessed into a form suitable for plotting reshaping data frames, deriving variables, or computing geometric relationships. Second, the visualization is assembled using ggplot2’s layered framework. These plots usually begin as script-level solutions that are often reused by copying both preprocessing and plotting code. Such cases often lead to their abstraction into utility functions [3]. Functions reduce errors, improve clarity, and ensure consistent behavior across analyses. The natural next step is to group related functions into a package. Even for personal or lab use, packaging brings clear benefits - organized code, internal documentation, and a consistent interface for future work. Sharing a package with the broader R community benefits both authors and users. Users may find the provided functions useful in their own work. At the same time, they may identify bugs, suggest improvements, or contribute enhancements. Even without external users, the formal requirements for public release - such as those imposed by CRAN - are valuable. Meeting these standards improves both the code and the skills of its authors.

Here, we introduce ggrecipes, an R package providing a collection of custom visualizations. These range from general-purpose plots, such as split-correlation, criteria, and rankshift plots, to domain-specific visualizations including sequence alignments, biodistribution plots, and kinetic rate (KD) maps. Each visualization serves as a basis that can be further customized using familiar ggplot2 syntax. Additional visualizations will be added in future releases. The package is available under an open-source MIT license. The source code and documentation are available at https://github.com/Ignophi/ggrecipes.

## 2. Methods

### Implementation

ggrecipes is organized around individual plotting functions that encapsulate both data preprocessing and visualization construction. The goal is to reduce preprocessing overhead while preserving ggplot2’s composability. Each function accepts a data frame and returns a ggplot2 object. Users can add additional geoms, modify scales, adjust themes, or apply coordinate transformations. Dependencies are kept to a minimum to reduce installation problems and potential conflicts. Beyond ggplot2, the package relies primarily on base R utilities and packages already imported by ggplot2 itself. Additional dependencies are limited to ggrepel [4] for non-overlapping label placement. Internal helpers for domain-specific data wrangling are not exported by default but can be accessed using the ::: operator.

### Code and data availability

All data used to demonstrate package functionality in this article consist of simulated example datasets that are included directly within the package vignettes and source code. Two real-world datasets, `mtcars`and `iris`, which are distributed with base R, are used as illustrative examples where appropriate. The complete code used to generate all figures shown in this manuscript is included in the released package, which is officially available on CRAN.

### Package development

Package development was carried out in R (version 4.4.1) using *devtools* [5] (version 2.4.6) and *usethis* [6] (version 3.2.1). The documentation website was generated with *pkgdown* [7] (version 2.1.3), and the package logo was created using *hexSticker* [8].

### Installation

The stable release of ggrecipes is available from the Comprehensive R Archive Network (CRAN) and can be installed in R using `install.packages(“ggrecipes”)`. The development version is publicly available on GitHub and can be installed using devtools: `devtools::install_github(“Ignophi/ggrecipes”)`. The CRAN package index and source repository are available at https://cran.r-project.org/web/packages/ggrecipes/index.html and https://github.com/Ignophi/ggrecipes,respectively.

## 3. Features

### 3.1 Split-correlation heatmap

Correlation heatmaps are widely used to visualize pairwise relationships between features. Many R packages provide functions for this purpose [9, 10, 11]. Standard correlation heatmaps are symmetric, displaying identical information in both triangles. When comparing two groups, `gg_splitcorr`uses this redundancy by displaying one group’s correlations in the upper triangle and the other group’s in the lower triangle.

The function requires a data frame with numeric variables and a binary splitting variable. It computes pairwise correlations for each group, adjusts p-values for multiple testing within each, and labels only statistically significant correlations with their values. **Figure 1** shows an example using the `mtcars`dataset split by engine type (V-shaped vs straight). The diagonal line separates the two groups, with labels indicating which group occupies each triangle. Correlation values are displayed only for significant pairs (p < 0.05 after adjustment).

**Figure 1.**
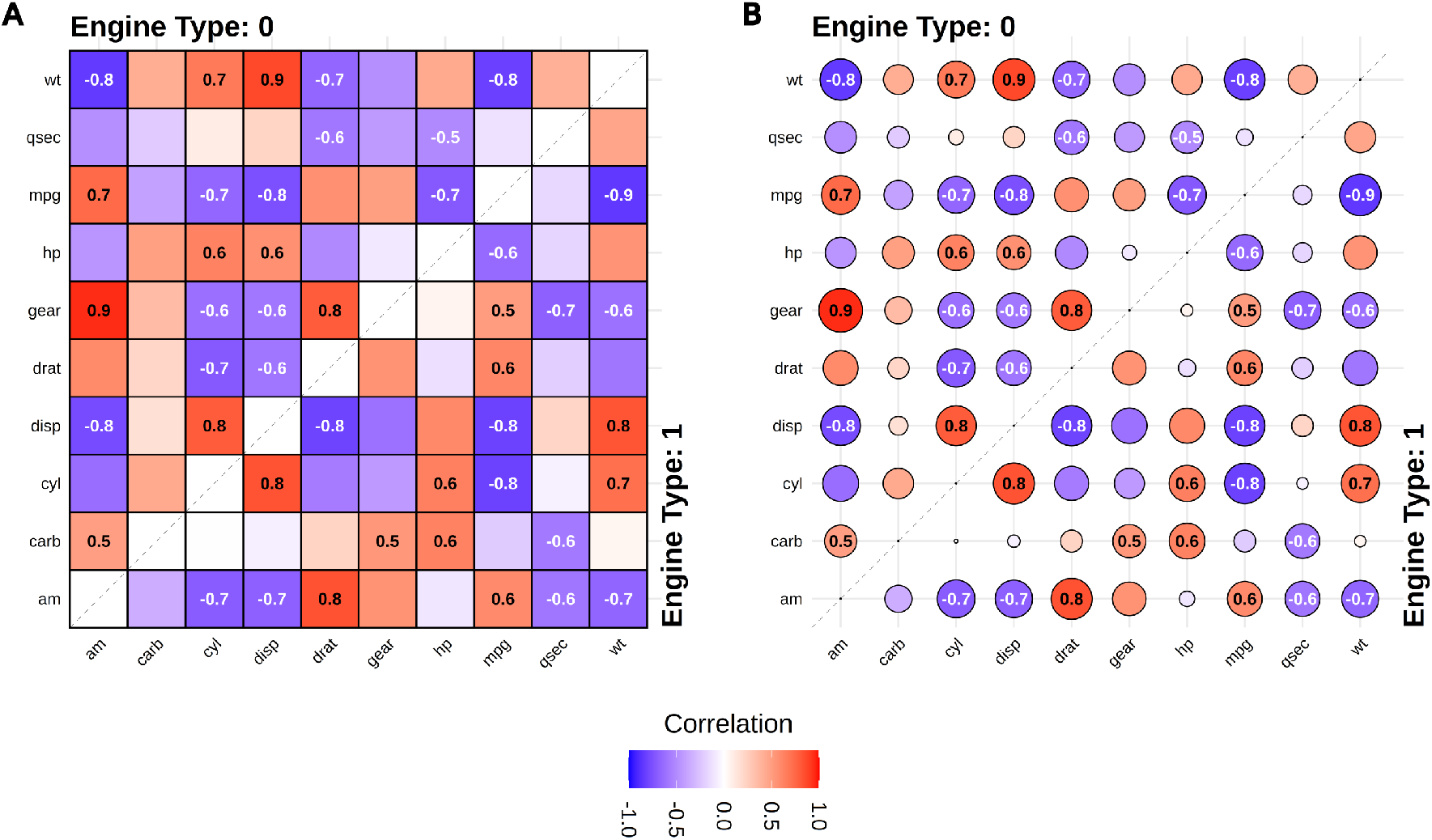
Split-correlation heatmap. Correlation analysis of the mtcars dataset split by engine type (0 = V-shaped, 1 = straight). The upper triangle displays correlations for engine type 0, while the lower triangle displays correlations for engine type 1. Only statistically significant correlations (p < 0.05 after Benjamini-Hochberg adjustment) are labeled with their values. Colors represent correlation strength from blue (negative) to red (positive). The diagonal line separates the two groups. **(A)** Tile-based visualization (style = “tile”). **(B)** Point-based visualization (style = “point”), where point size indicates absolute correlation magnitude.

The visualization style can be switched from tiles to points using the style argument **(Figure 1B)**. Point size then represents absolute correlation magnitude. Users can customize correlation methods (Pearson or Spearman), p-value adjustment procedures, color schemes, and aesthetic details.

### 3.2 Rank shift plots

A frequent question concerns changes in sample ranking between two conditions. `gg_rankshift`creates a three-panel visualization: side panels display ranked distributions for each condition, while the center panel connects corresponding samples with lines colored by rank change direction. The function requires data with sample identifiers, a grouping variable containing exactly two levels, and a numeric value for ranking. Ranks are calculated using summary statistics (mean or median) controlled by `stat_summary`. Distributions can be displayed as barplots **(Figure 2A)** or boxplots **(Figure 2B)** via the style argument. Individual measurements can be overlaid using `show_points`to display replicate variability. The `free_x`argument controls whether side panels share x-axis limits or use independent scales. Panel widths are controlled by `panel_ratio`, while colors are customizable through `fill_colors`for distributions and `rank_change_colors`for connecting lines.

**Figure 2.**
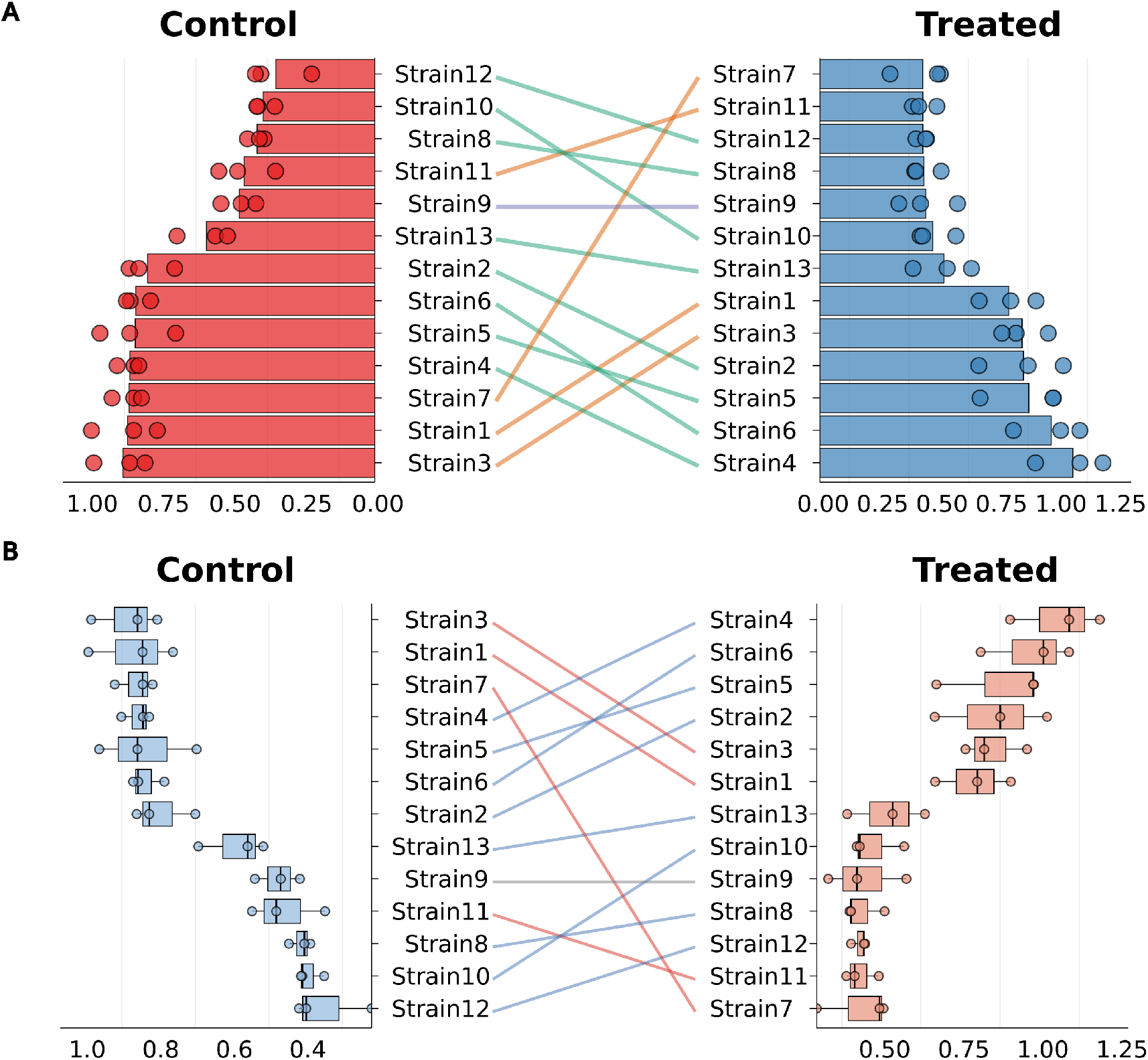
Rank shift plots. Changes in sample ranking between control and treated conditions for 13 strains. The left and right panels show ranked distributions for the control and treated groups, respectively, while the center panel connects corresponding strains across conditions. Lines are colored by rank change direction, indicating upward shifts (blue), downward shifts (red), or no change (grey) in rank between conditions. **(A)** Barplot-based visualization, where bars represent summary statistics (mean) for each strain and individual measurements are overlaid as points. **(B)** Boxplot-based visualization showing the distribution of replicate measurements for each strain. In both panels, strains are ordered independently within each condition based on their rank, highlighting reordering between conditions rather than absolute values.

### 3.3 Criteria plots

In many settings, samples are compared across several categorical criteria [12]. `gg_criteria`displays this information as a heatmap with samples on the y-axis and criteria on the x-axis. Criterion columns are se-lected by a regular-expression pattern and converted to a long format internally. Each tile encodes the categorical value (e.g., Yes/No, residue identity) using discrete colors, with optional text labels, while preserving the original sample order. **Figure 3** illustrates gene prioritization based on multiple genetic and functional observations, with a single “Yes” level shown for each criterion. When one or more columns are supplied via `bar_column`, the function adds horizontal bar plots whose rows align with the heatmap. Panel widths are controlled by `panel_ratio`, and bar colors can be customized or drawn from a Brewer palette. **Figure 3** uses a single barplot to show the total number of supporting criteria per gene, while **Figure 4** shows three metrics (binding affinity, expression yield, and thermal stability) for a panel of mutants evaluated against positional mutation criteria. As the vertical alignment between heatmap and barplot panels depends on the exported figure dimensions, the function provides recommended sizes based on the number of samples and criteria.

**Figure 3.**
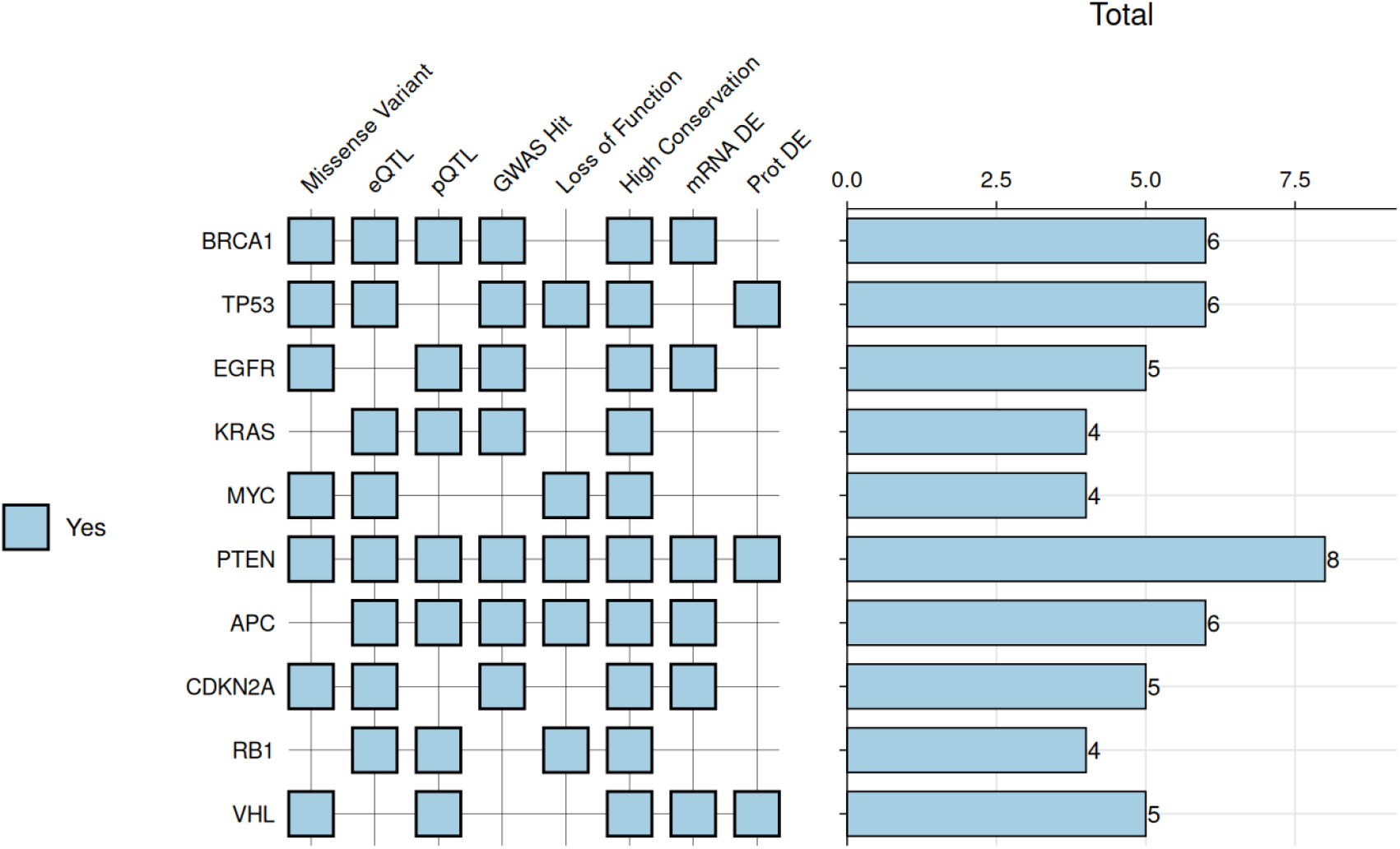
Criteria plots - gene prioritization. Visualization of multiple categorical criteria across a set of genes. Rows correspond to genes and columns correspond to individual criteria derived from genetic, functional, and expression-related observations. Each tile indicates whether a given gene satisfies a specific criterion, with filled tiles denoting a positive (“Yes”) assignment. The right panel shows a horizontally aligned bar plot summarizing the total number of criteria satisfied by each gene.

**Figure 4.**
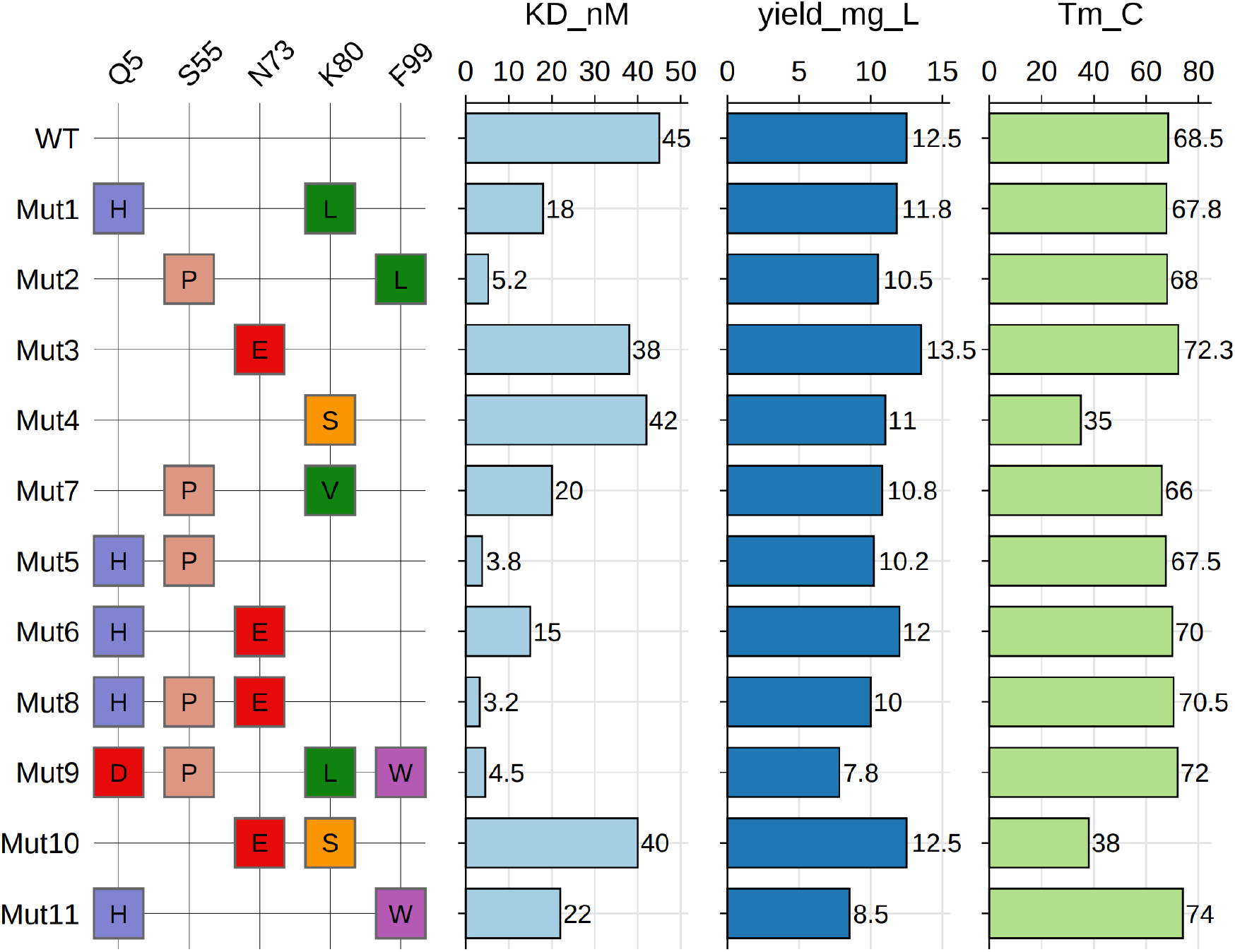
Criteria plots - protein mutant characterization. Example of a criteria plot generated using the same function as in Figure 3, illustrating its flexibility for more complex layouts. The criteria panel displays positional mutation information for a set of protein variants, with rows corresponding to variants and columns corresponding to mutation sites. Tiles indicate the substituted residue at each position, and only mutated positions are shown. Additional horizontally aligned bar plot panels are included to display quantitative measurements for each variant, including binding affinity (KD, nM), expression yield (mg/L), and thermal stability (Tm, °C).

### 3.4 Genotype plots

Genotype data typically consist of biallelic markers such as SNPs, where each individual carries two alleles at each locus. `gg_geno`provides a specialized visualization for these data, sharing the core design and features of `gg_criteria`but adapted for diploid genotypes. Each tile is split diagonally to represent both alleles simultaneously, with samples occupying rows and genetic markers occupying columns.

The function accepts data in wide format with genotype columns identified by a regular expression pattern. Genotypes must be encoded as “allele1/allele2” for unphased data or “allele1|allele2” for phased data. The function automatically detects phasing based on the separator and visually distinguishes the two cases using border colors: black borders indicate phased genotypes, while white borders indicate unphased genotypes. The top-left triangle displays the first allele, and the bottom-right triangle displays the second allele. **Figure 5** shows genotype data for twelve individuals across six SNPs. Alleles are color-coded (0, 1, 2, 3), with heterozygous genotypes immediately visible as split-colored tiles.

**Figure 5.**
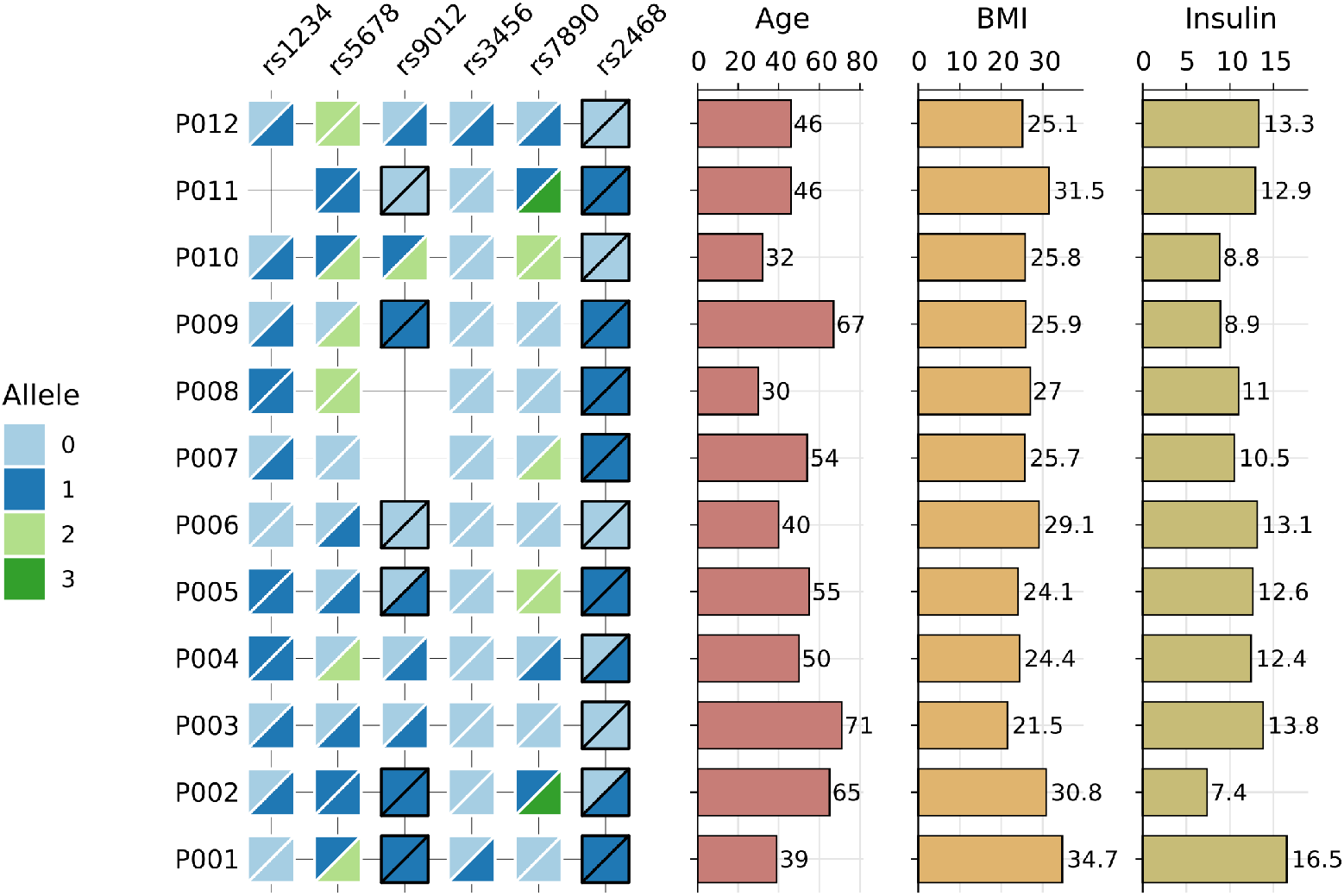
Genotype plots with aligned phenotypic measurements. Visualization of diploid genotype data across multiple single-nucleotide polymorphisms (SNPs) for a cohort of individuals. Rows correspond to individuals and columns correspond to SNP loci. Each tile is split diagonally to represent the two alleles carried by an individual at a given locus, with colors indicating allele identity. Phased genotypes are indicated by black tile borders, while unphased genotypes are indicated by white borders. Additional horizontally aligned bar plot panels display continuous phenotypic variables for each individual, including age, body mass index (BMI), and insulin levels.

Like `gg_criteria`, the function supports horizontal barplots via `bar_column`to display continuous variables alongside genotypes. **Figure 5** includes three barplots showing age, body mass index (BMI), and insulin levels for each individual, enabling direct comparison of genetic variation and associated phenotypes. The function shares customization options with `gg_criteria`, including `panel_ratio`for barplot width, `tile_width`and `tile_height`for tile dimensions, `border_width`for tile borders, and `tile_fill`for custom allele colors.

### 3.5 Sequence plots

Visualizing sequence mappings to a reference is a common task in proteomics and genomics, arising in peptide mapping [13], read alignment, and variant analysis. `gg_seq`displays where sequences match within a reference, with each sequence occupying a horizontal row ordered by starting position. **Figure 6** shows peptide mapping of an antibody fragment. Specific amino acids can be emphasized with custom colors using the color argument. For example, lysines and arginines can be colored to indicate trypsin cleavage sites. The highlight argument adds vertical shading bands at specified positions, useful for marking regions such as CDRs, mutation sites, or functional domains. The `name`argument displays sequence identifiers on the y-axis. The `wrap`argument allows users to specify the number of positions to display per row before wrapping to the next panel. The `annotate`argument adds text labels above the plot with customizable positioning and styling. The reference sequence can be shown or hidden with `show_ref`.

**Figure 6.**
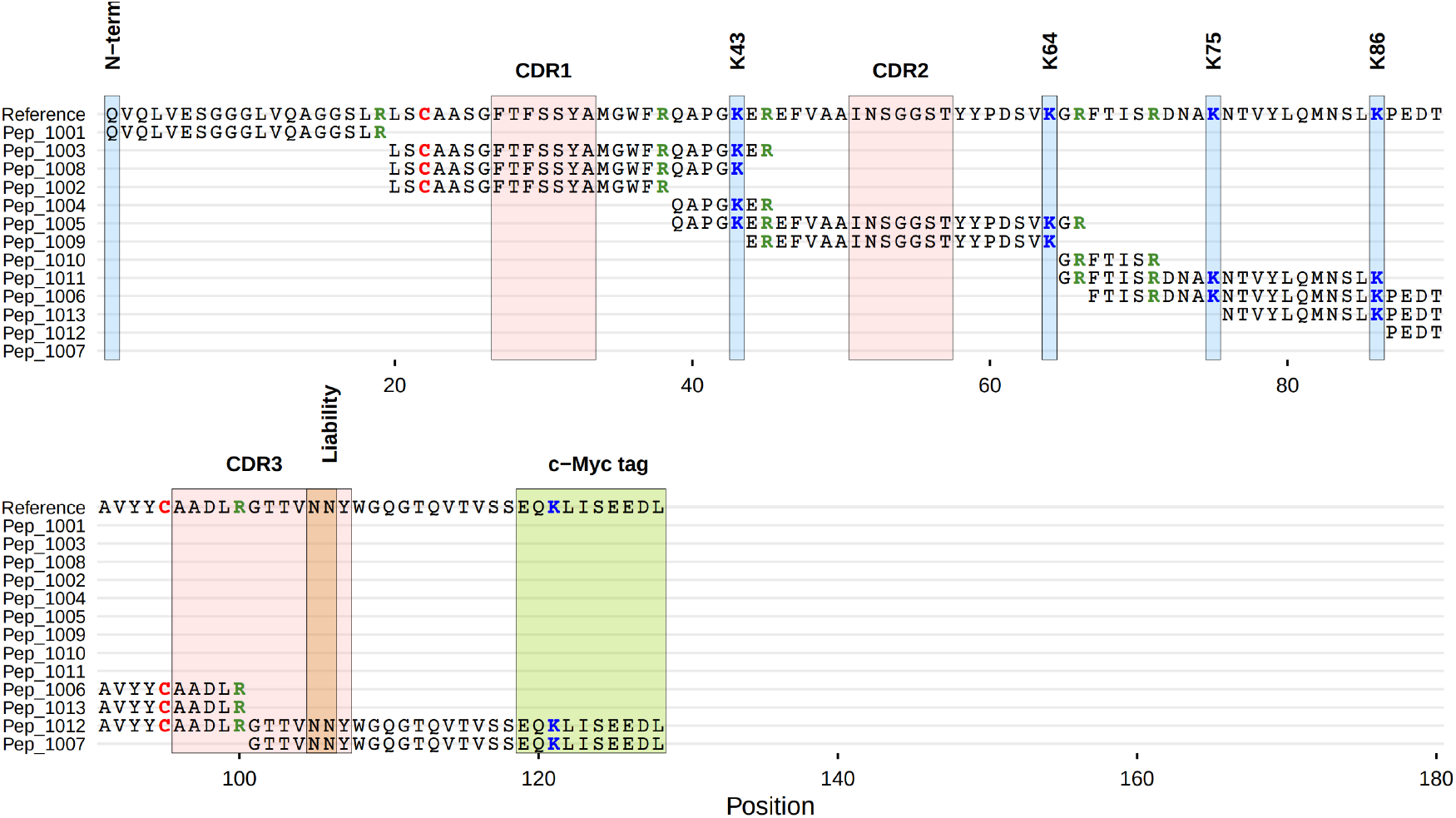
Sequence plots - Peptide mapping coverage. Visualization of peptide coverage along a reference protein sequence. Rows correspond to individual peptides and columns correspond to positions along the reference. Characters indicate amino acid identities at covered positions, while uncovered regions are left blank. Peptides are ordered by their starting position along the reference sequence. Vertical shaded bands highlight annotated regions and positions of interest, including N-terminal residues, complementarity-determining regions (CDR1–3), selected lysine positions, and a C-terminal c-Myc tag. Amino acids can be colorcoded to emphasize specific residue types. Position indices are shown on the x-axis.

For sequence variant analysis, a common approach is displaying only substituted positions to reduce visual noise (e.g., Jalview’s [14] “Show Differences from Reference” option or MSA viewers with consensus masking). `gg_seqdiff`displays only positions that differ from the reference, with matching positions hidden. Only differing characters are displayed, making mutations immediately visible **(Figure 7)**. The function can also parse Clustal alignment files directly using the `clustal`argument. `gg_seqdiff`supports the same customization options as `gg_seq`.

**Figure 7.**
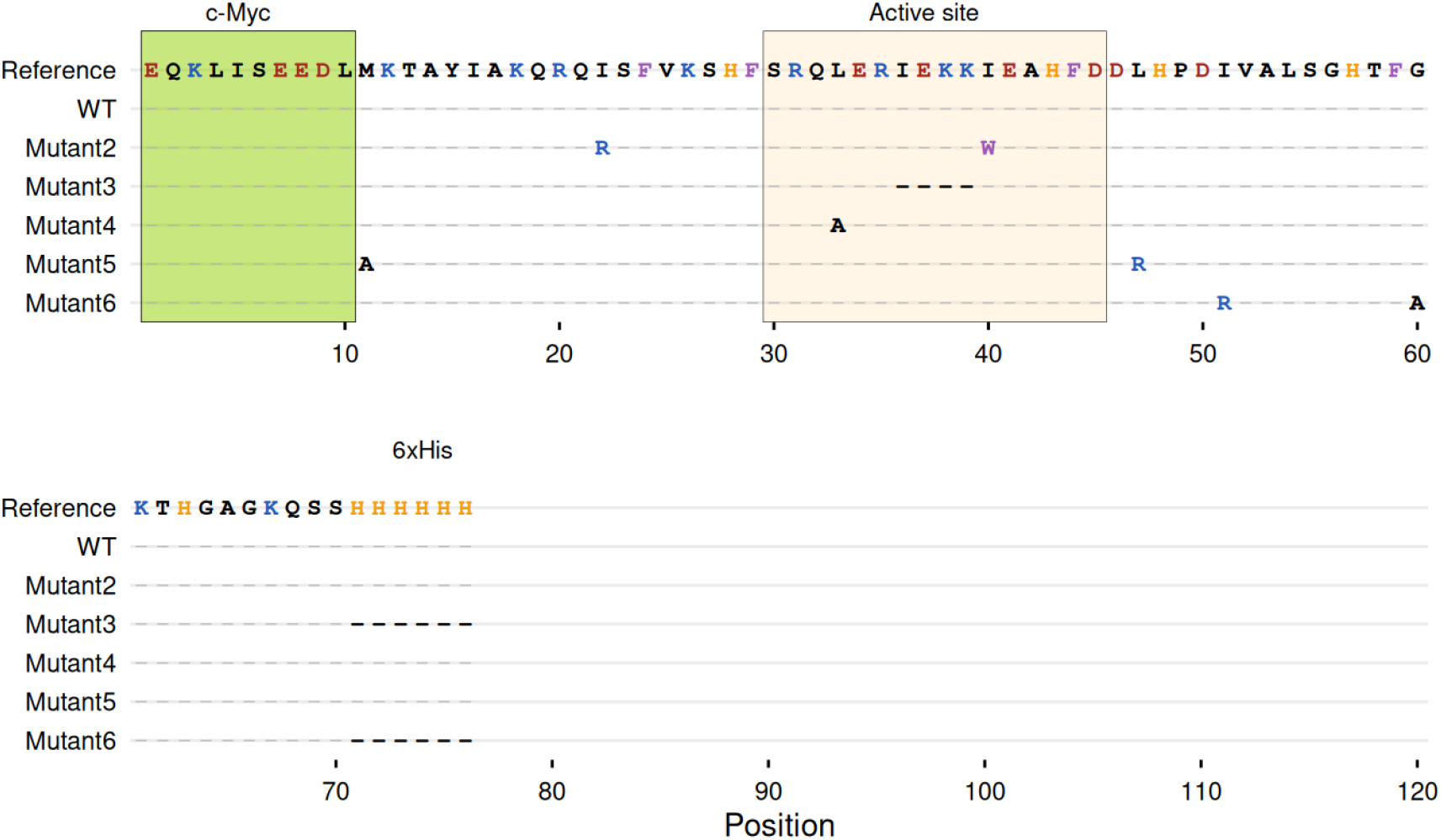
Sequence difference plots. Visualization of sequence variants relative to a reference sequence, showing only positions that differ from the reference. Rows correspond to individual variants and columns correspond to positions along the reference. Identical positions are omitted, while substituted residues are displayed at their corresponding locations, reducing visual clutter and emphasizing sequence differences. Vertical shaded bands highlight annotated regions and positions of interest, including a c-Myc region, an active site, and a C-terminal 6×His tag. The reference sequence is shown for context, and position indices are shown on the x-axis.

### 3.6 Biodistribution plots

Biodistribution studies assess how constructs distribute across organs and tissues, commonly reported as percent injected dose per gram (%ID/g) or similar metrics. Visualizing these data presents a challenge when uptake varies widely across organs - high-uptake organs such as liver or kidney can compress the scale, making differences in low-uptake organs difficult to discern. One approach is to use y-axis breaks, but this becomes unwieldy with multiple high-uptake organs and high variability, and requires case-by-case adjustment. `gg_biodist`addresses this through faceting, allowing specific organs to be displayed on independent y-axis scales while maintaining a unified visual presentation.

The function accepts data in either long or wide format. For wide format, measurement columns are identified by a regular expression pattern. **Figure 8A** shows biodistribution of an antibody-drug conjugate across multiple organs. Bars represent mean values, with individual measurements overlaid as points. The separate argument accepts a vector of organ names to display on independent y-axis scales. **Figure 8B** demonstrates this with tumor and kidney uptake separated into distinct facets. The function supports grouped comparisons through the group argument. The `fill_colors`argument accepts custom color vectors, with length matching the number of groups, defaulting to a Brewer palette for grouped data or single blue color otherwise. Error bars displaying standard deviation can be added via `error_bars`.

**Figure 8.**
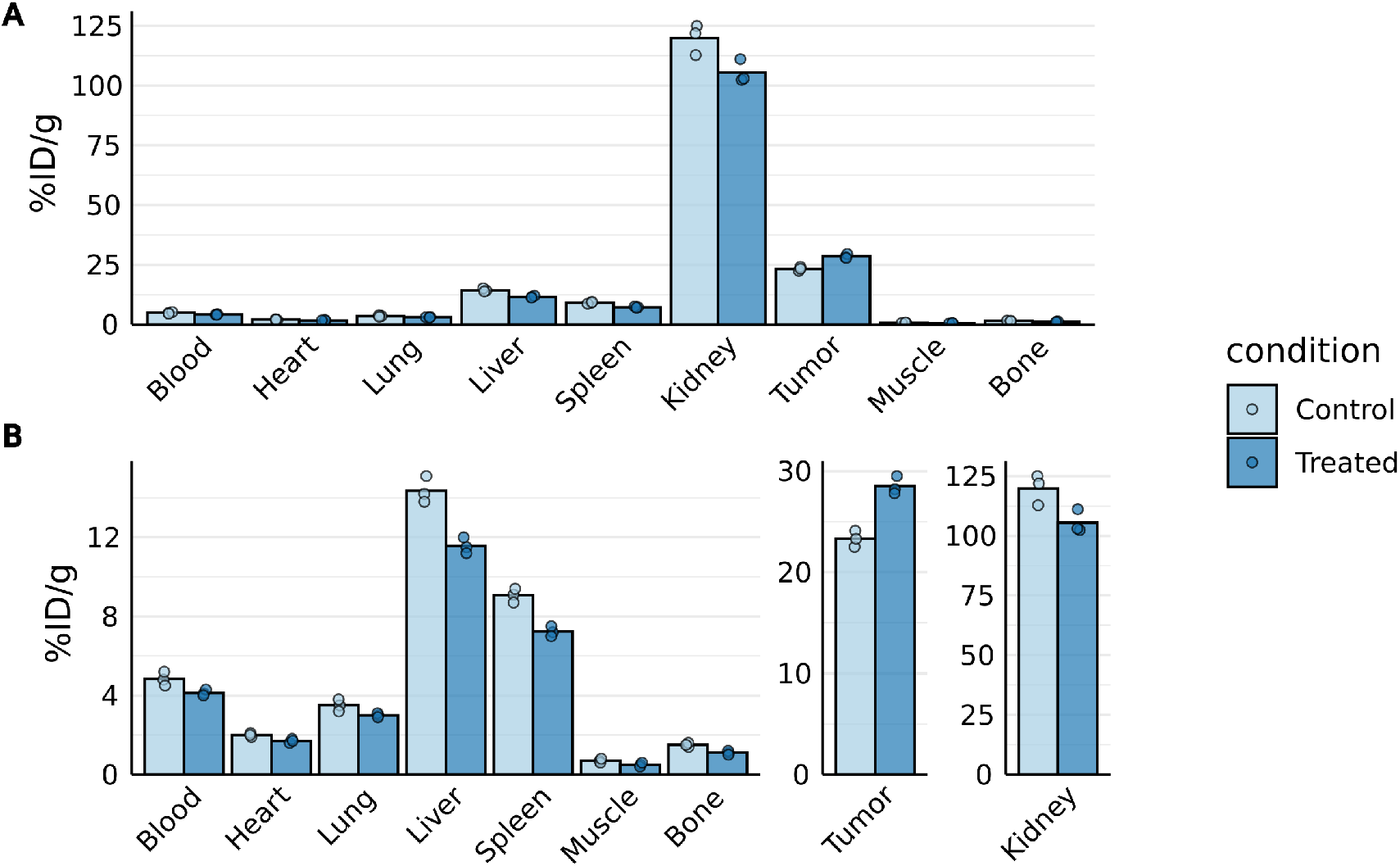
Biodistribution plots. Visualization of biodistribution measurements (% injected dose per gram, %ID/g) across multiple organs under control and treated conditions. Bars represent mean values, with individual measurements overlaid as points. **(A)** Standard grouped bar plot displaying all organs on a shared y-axis scale, where high-uptake organs dominate the dynamic range. **(B)** Faceted visualization in which selected organs are displayed on independent y-axis scales while maintaining a unified visual presentation. This approach enables clearer inspection of lower-uptake organs without isolating panels or breaking alignment across the figure. Tumor and kidney are shown on separate scales.

### 3.7 Kinetic rate maps

Kinetic binding data from assays such as surface plasmon resonance (SPR) [15] or biolayer interferometry (BLI) are often summarized as association (ka) and dissociation (kd) rate constants. `gg_kdmap`displays these measurements on a log-log plot with kd on the x-axis and ka on the y-axis. Diagonal iso-affinity contours indicate constant equilibrium dissociation constants (KD = kd/ka), so points along the same line share the same “affinity”. This allows one to understand visually whether similar affinities arise from faster association, slower dissociation, or a combination of both. The function requires a data frame with columns for association rate (ka, in *M*^−1^*s*^−1^), dissociation rate (kd, in *s*^−1^), and an identifier for grouping replicates. **Figure 9** shows kinetic data for antibody variants: each point represents a measurement, with replicate measurements connected by lines. Diagonal dashed lines indicate constant KD values, which are automatically calculated and labeled on secondary axes in appropriate units (pM, nM, µM, or mM) based on the data range. A reference such as a wild-type or benchmark molecule can be highlighted via `ref_id`. The `show_anno`argument adds text annotations indicating fast on-rate and fast off-rate regions. The labels argument displays identifiers using *ggrepel* for non-overlapping placement. Iso-KD lines are spaced evenly across the visible KD range in log space. These can be customized using various arguments.

**Figure 9.**
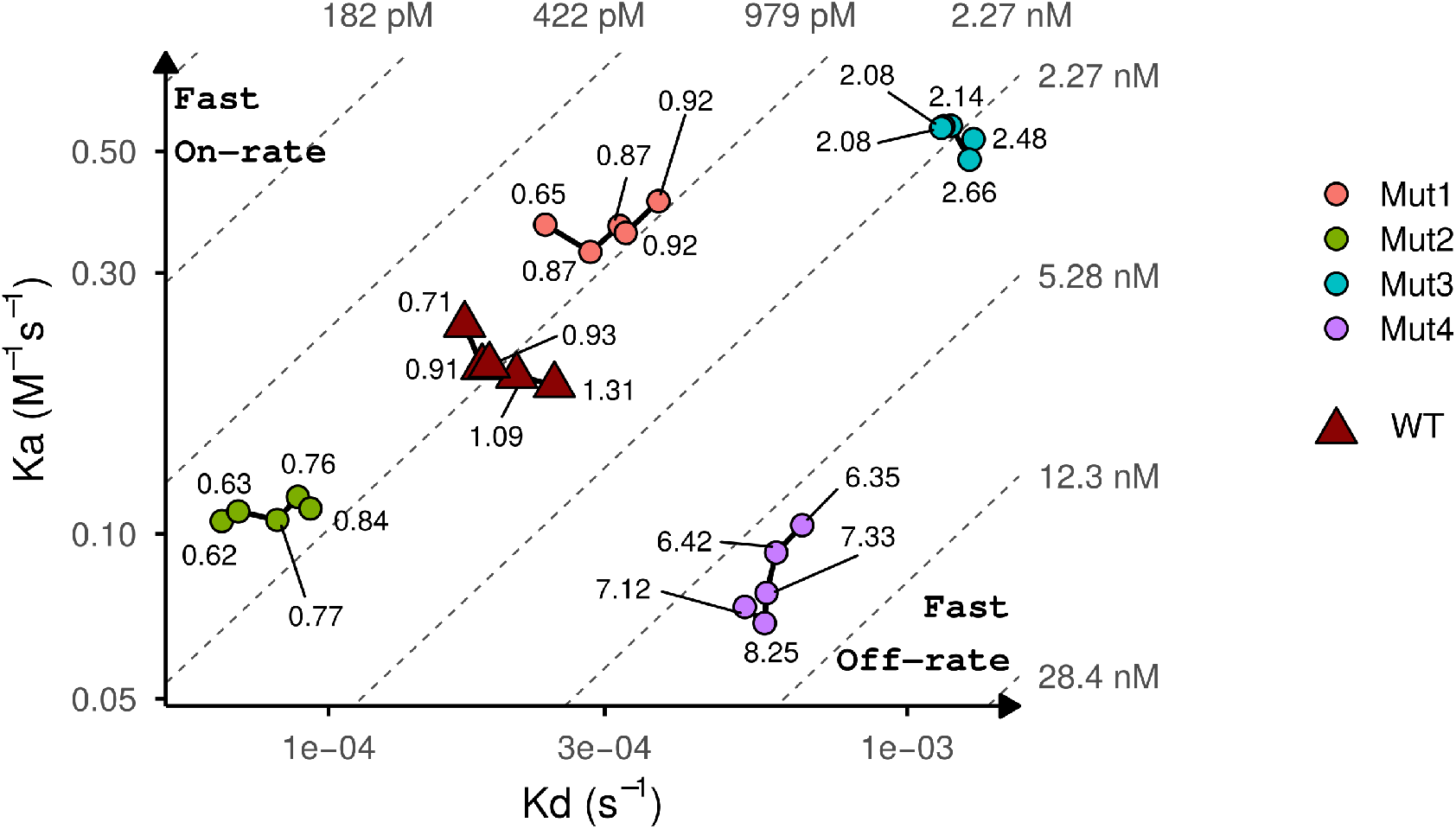
Kinetic rate maps. Visualization of kinetic binding parameters on a log–log scale, with dissociation rate constants (kd, *s*^−1^) on the x-axis and association rate constants (ka, *M*^−1^*s*^−1^) on the y-axis. Each point represents a kinetic measurement, with replicate measurements for the same variant connected by lines. Diagonal dashed lines indicate constant equilibrium dissociation constants (KD = kd/ka), with corresponding KD values labeled in appropriate units. Points lying along the same diagonal share the same affinity but differ in their association and dissociation kinetics. The wild-type reference is shown separately from variant groups. Annotations indicate regions corresponding to fast on-rates and fast off-rates.

## Acknowledgments

The book “R Packages” [3] was a great help in the development of this package. I would like to thank the authors and maintainers of *devtools, usethis*, and *pkg-down*, which substantially simplified package development and documentation.

## Correspondence

If you find any mistakes or have certain comments, I would appreciate it if you can let me know at.

